# The Effects of Living at Altitude on the Prevalence of Dementia

**DOI:** 10.1101/2025.01.03.630371

**Authors:** Alan Holt, Adrian Davies

## Abstract

A lower incidence of dementia has been observed in populations residing at high altitudes compared to those at low altitudes. A possible cause is the inverse relationship between altitude and the partial pressure of inspired oxygen. This paper investigates the long term impact that living at altitude has on mitochondrial function.

We use simulation methods to model the dynamics of a population of mtDNA and the proliferation of deletion mutants. Clonal expansion of these deletion mutants over many years impairs mitochondrial function resulting in cell death, notably in post-mitotic cells such as neurons. Neuron loss leads to cognitive impairment, progressing to dementia over time.

Our results show that there is a moderate reduction in neuron loss rates and associated cognitive decline with respect to dementia in the elderly. These results are consistent with published data on the regional prevalence of dementia when plotted against altitude.

## 1 Introduction

In this paper we study the long term effects of living at altitude on the prevelance of dementia. Specifically, we focus on mitochondrial dysfunction caused by the accumulation of mtDNA deletion mutants (mtDNA_*del*_) over a human lifespan. Dementia is a symptom of a number of pathologies but it is reasonable to assume that there are factors in common amongst these pathologies that result in neuron loss and consequent dementia [3, 2]. A plausible candidate is mitochondrial dysfunction [10]. Neuron loss can ensue from the proliferation of mtDNA_*del*_ out competing the wild-type (mtDNA_*wild*_) resulting in impaired mitochondrial function. Neuron loss rates beyond those seen in normal aging can lead to dementia of varying degrees.

At high altitude, oxygen levels are lower due to the decreased atmospheric pressure. The absolute percentage of oxygen in the atmosphere remains at 21% regardless of altitude, however, with an increase in altitude the partial pressure of oxygen and the amount available for metabolism, decreases. Chronic exposure can lead to hypoxia, which can be detrimental to brain health [26]. Mild hypoxia, however, may stimulate the production of neuro-protective factors and enhance cognitive function [4]. Living at high altitudes can also affect cardiovascular health which is closely linked to the risk of developing dementia [19].

Transitioning to a high altitude results in a short term increase in blood pressure and heart rate [32, 17]. Midlife hypertension, diastolic pressure and a high resting heart rate are associated with increased risk of dementia [17, 28]. These same symptoms can be experienced by someone that moves to high altitude, however, the symptoms are temporary as the body acclimatises. People who live at high altitudes may have different lifestyles and environmental exposures compared to those at lower altitudes. Factors, such as diet, physical activity and sun exposure can all play a role in cognitive health [8]. Long term adaptation to the decreased partial pressure of oxygen involves various metabolic processes including cellular respiration [25].

Dementia is a complex condition which is likely influenced by multiple factors. While some studies have explored the effects of altitude on cognitive function and brain health, the results are mixed and often inconclusive [20, 12, 37]. We modeled a specific aspect of the effect of reducing the partial pressure of oxygen and the effect that this would have on mtDNA in neurons consistent with living at altitude. Altitude may play a role in the prevalence of dementia, however, it is likely to be just one of many contributing factors. Nevertheless, we were curious as to how living at altitude would affect mtDNA dynamics within our simulated organelle and the resultant levels of cognitive dysfunction due to neuron loss. We ran simulations for hosts with a high predisposition towards dementia at altitudes of 1,600 and 3,000m. We compared these results to hosts living at sea level with low and high predisposition. We observed the effect on mtDNA_*del*_ proliferation within mitochondria and consequent neuron longevity. We determine neuron loss rates and assess the severity of cognitive dysfunction, specifically, with regard to dementia in the elderly. Neuron loss rates associated with dementia, both global and local, are disputed. As such, there is little consensus in the literature as to the rates of neuron loss that constitute dementia [11, 7, 33, 2]. For the purposes of this paper we select a target of 20-40%, where 20% is the onset of dementia and 40% is advanced. Below 20%, we consider normal aging but above 40% it is likely the host would not have survived.

The contribution of this paper with regard to the long term effects of living at altitude on the prevalence of dementia, is three-fold:

- Presents an analysis in order to establish a link between living at altitude and the prevalence of dementia.
- Identifies how altitude would effect the parameters in our simulation of mtDNA proliferation and to what degree.
- Shows that, for a host with a high predisposition to dementia, living at altitude would delay onset by a few years and would yield a moderate reduction in severity.

### 2 Altitude and Dementia

In this section we review the link between living at altitude and prevalence of dementia. A meta-analysis by Urrunaga-Pastor *et al*. [31] reports an *increase* in cognitive dysfunction in the elderly at altitude. However, the paper advises caution in interpreting the results. Given that the respective studies analysed by Urrunaga-Pastor *et al*. were from different countries, heterogeneity in clinical assessment of cognitive dysfunction was to be expected.

In Fig. 1 we plotted dementia rates (as a percentage of the population) of a number of European countries [1] against mean altitude for the respective country [35]. It can be seen that there is a downward trend, however, confidence intervals (translucent band) are quite broad suggesting a lack of statistical significance. As with the analysis presented in Urrunaga-Pastor *et al*. there is a problem of methodological heterogeneity given that the data is compiled from different health jurisdictions. Furthermore, there is not a wide range of mean altitudes between European counties, where the lowest is 30m (Netherlands) and the highest is 1,132m (Turkey).

**Figure 1:**
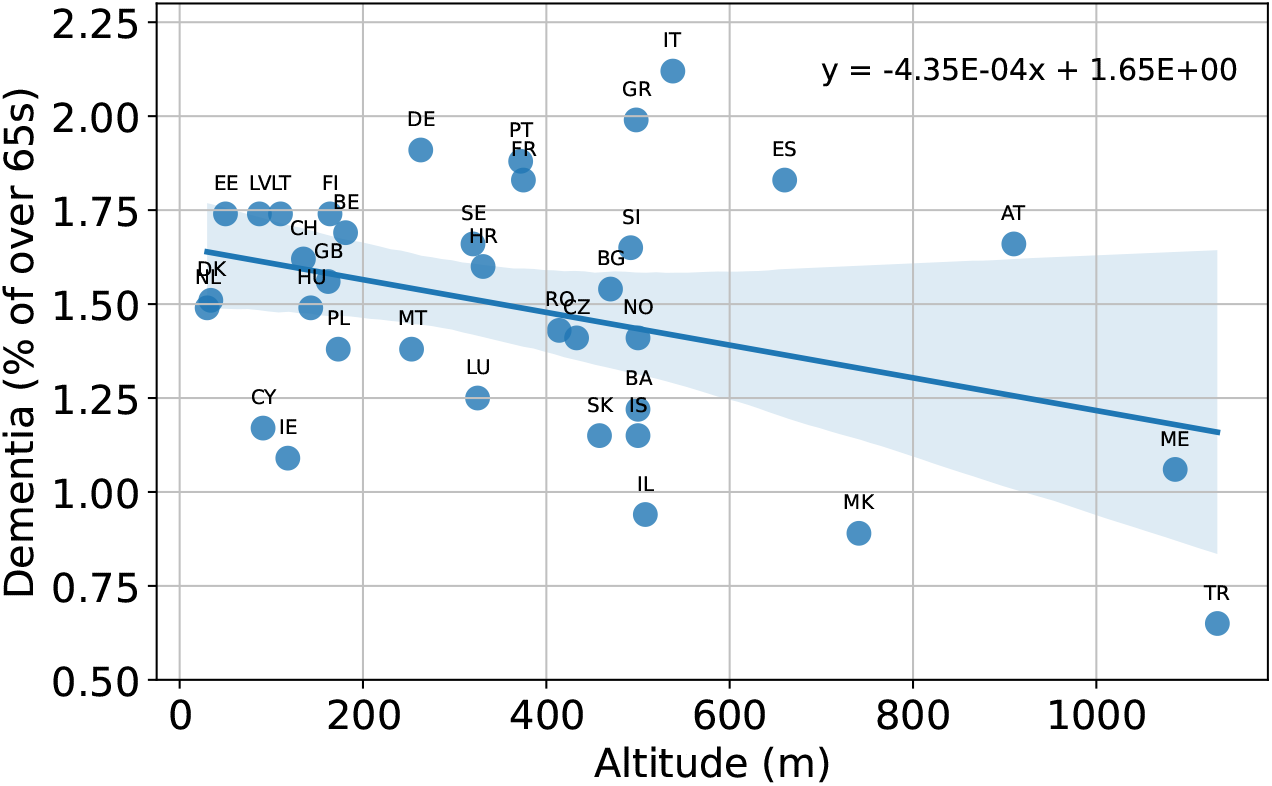
European countries: percentage of population with dementia versus altitude (shaded area indicates error bars).

The graph in Fig. 2 shows dementia rates for Chinese provinces presented in Wu *et al*. [36] versus altitude [35]. As with the European data, the confidence intervals are quite broad (blue). This is due to the Sichuan data point which could be an outlier. Omitting it, reduces the confidence interval range of the resultant regression model (orange). We appreciate that the practice of removing inconvenient data points may be questionable, however, the percentage of the population over 65 is much higher in Sichuan province than the national average [9, 23], which may explain its outlier status. The advantage of the Chinese province data set over that of the European country data is that, it comprises data points for higher altitudes (the highest province being 3,710m). Furthermore, the paper claims to use the Diagnostic and Statistical Manual of Mental Disorders (DSM) which obviates methodological heterogeneity.

**Figure 2:**
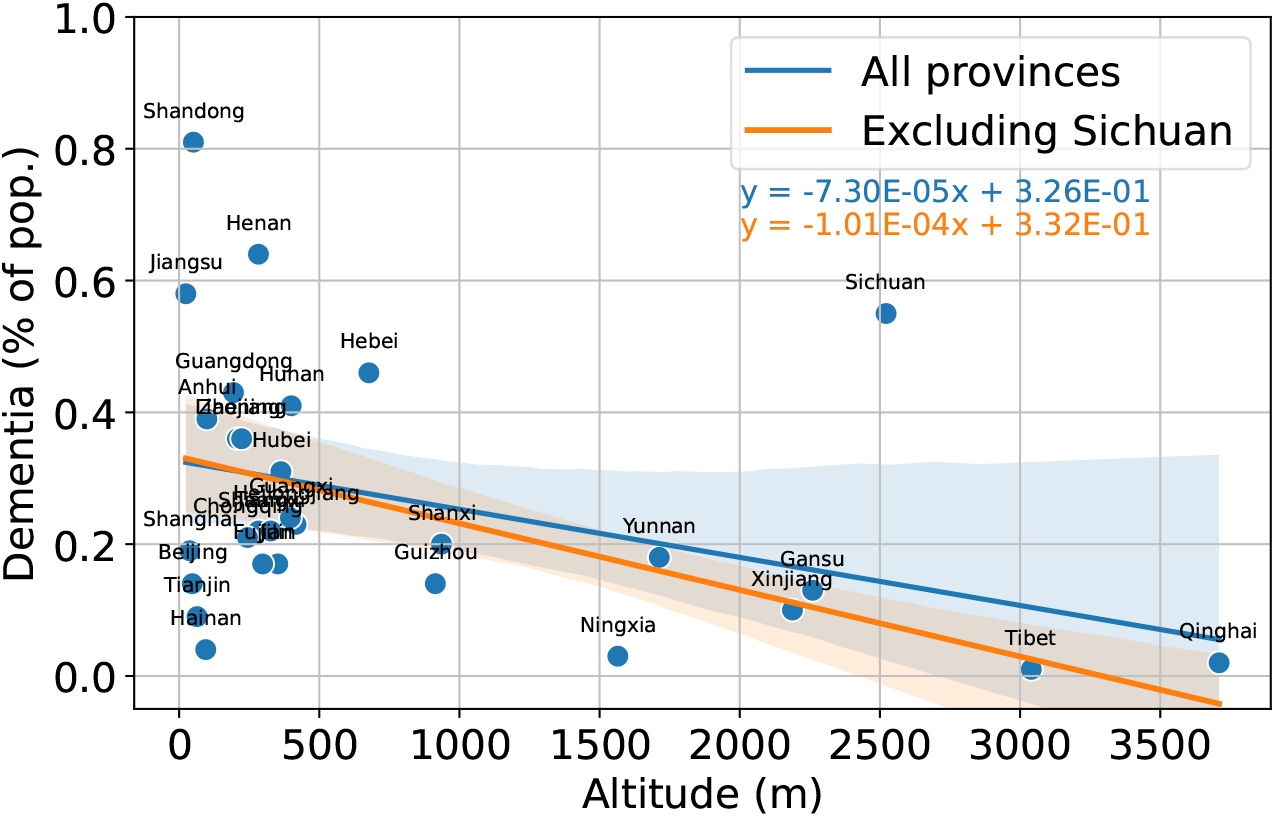
Chinese Provinces: Percentage of population with dementia versus altitude.

In our final analysis we examine per-county data from the United States which accompanies the study by Dhana *et al*. [6]. Using US counties overcomes some of the limitations of the analyses described above:

- The US is a large geographical area with over 3,000 counties across 50 states.
- Dhana *et al*. [6] adopts DSM. item There is a broad range of altitudes, where the average per-county ranges from sea level to over 3,600m.
- Some studies of dementia versus altitude exist for certain parts of the US, namely, California [30].

County altitudes were surprisingly difficult to come by. After an exhaustive search we were unable to find a source of batch data for US county altitudes. Values were available interactively from sites such as [35] but making over 3,000 separate interactive requests would have been prohibitively time consuming. We resorted to writing a script to perform repeated Google searches. With this method we were able to retrieve data for most counties which left a manageable amount that could be looked up interactively. The per-county dementia versus altitude data is shown in Fig. 3. There is a small decrease in dementia prevalence with altitude with low range of confidence interval. We highlight the dementia results in Fig. 3 for the state of California. These results align with the those presented in Thielke *et al*. [30] which showed that, across the counties of California, there was a decrease in the mortaity due to Alzheimer’s with altitude.

**Figure 3:**
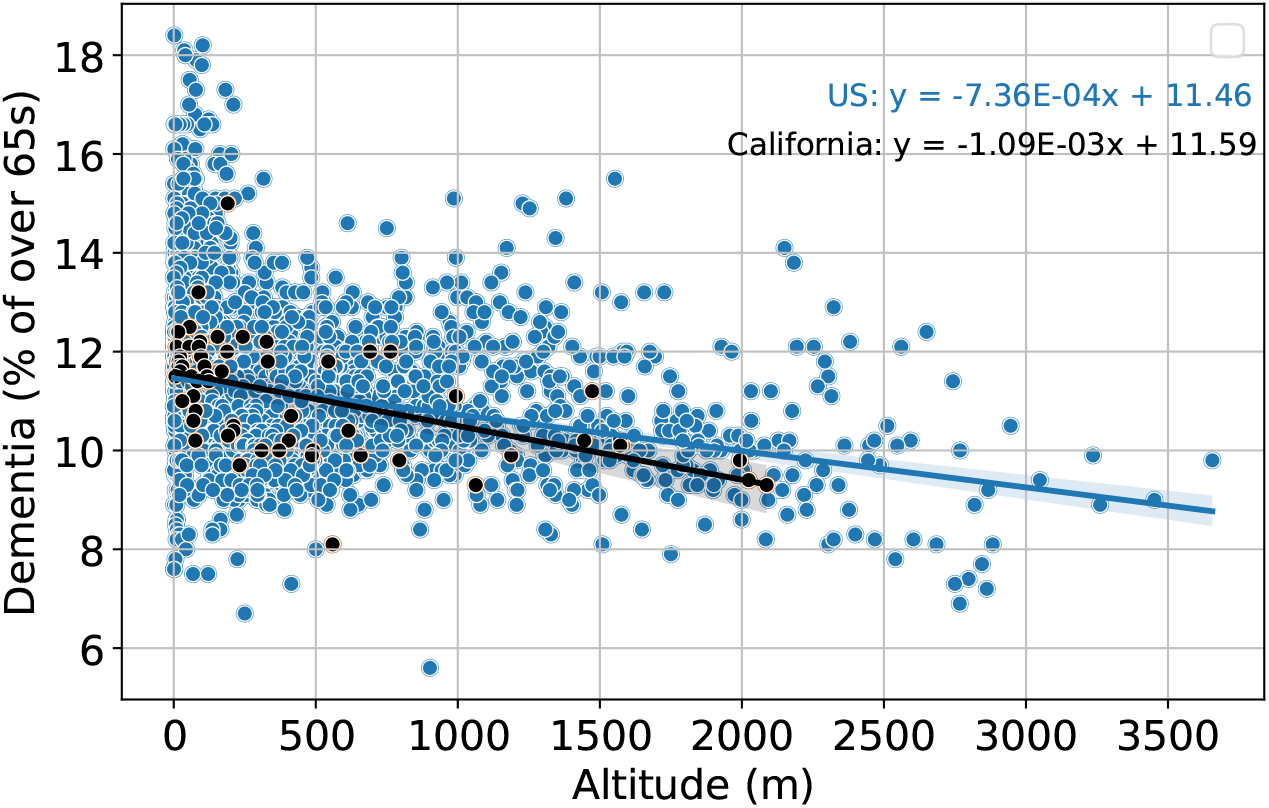
Prevalence of dementia versus mean US county altitude.

From the analyses above, it is difficult to establish a definitive relationship between dementia prevalence and living at altitude. Elevations will fluctuate considerably within country/province/county and there will be mobility in certain sections of the population. Dementia is a symptom of many different pathologies which exacerbates the problem of isolating any pattern as a result of living at altitude. However, despite the noise in the data, there does appear to be a downward trend in dementia prevalence with altitude. From these results, we are inclined to believe, living at altitude would yield a moderate benefit with regard to the prevalence of dementia. Thus, we would expect a small reduction in neuron loss rates from our simulation results for hosts living at altitude compared to sea level.

### 3 Simulator

In this paper we simulate the proliferation of mtDNA_*del*_ amongst a population of mtDNA_*wild*_ within a pseudo mitochondrial organelle of a neuron. Our simulation is based upon the Mitochondrial Free Radical Theory of Aging (MFRTA). In our simulation mtDNA are software data structures that, like their biological counterparts, are subject to *expiration, replication, mutation* and *competition*.

The simulation runs in discrete time intervals of *t* = 15 minutes for a 100 year (human) lifespan whereby mtDNA may expire, replicate and/or mutate. In each *t* we record the state of the organelle in terms of the number of mtDNA in each species. While there is only one mtDNA_*wild*_ species there may be multiple species of mtDNA_*del*_.

Random events occur within the simulation according a Bernoulli trial success/failure for some probability, for example:

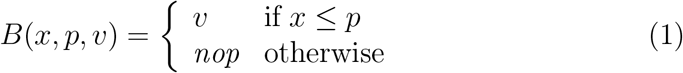

where 0 ≤ *x* ≤ 1 is a uniformly distributed random number, 0 ≤ *p* ≤ 1 is some probability threshold and *v* is the event. In our simulation, a success will result in the execution of some code that implements *v* and a failure results in a *nop* (no operation).

We implement a mechanism that expires mtDNA over time. Each mtDNA is given a time-to-live (TTL) value which is randomly decremented each interval according to the Bernoulli trial:

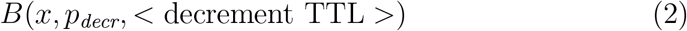

When the TTL reaches zero, the mtDNA expires and is removed from the population. The TTL and 0 ≤ *p*_*decr*_ ≤ 1 are calibrated to yield a 30 day half-life amongst all mtDNA.

Neurons need to maintain a population of mtDNA_*wild*_ despite mtDNA having a lifespan of several orders of magnitude shorter than the neuron itself. Being post-mitotic, a neuron, ideally, needs to last for the lifespan of the host. Maintenance of an mtDNA population is critical to the survival of the neuron as mtDNA_*wild*_ are responsible for energy production within it. Autophagy results from a depletion of the wild-type population as it is not able to meet the neuron’s energy demands. However, given mtDNA have a shorter lifespan than neurons, mtDNA must replicate in order to maintain this population. In our simulation, the opportunity to replicate is granted to an mtDNA upon success of the Bernoulli trial:

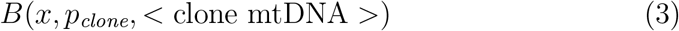

where *p*_*clone*_ = 0.01 *h*^−1^, which is pursuant to the mitigation of attrition due to Eq (2). However, mtDNA replication is subject to certain restrictions beyond the success of the Bernoulli trial, namely:

- The neuron has an energy deficit.
- The population of mtDNA does not exceed the organelle’s capacity *C*_*max*_.
- The mtDNA is not currently undergoing cloning.

All of the above need to be true, otherwise, the mtDNA cannot clone.

In order to prevent mtDNA_*wild*_ replicating without bound, we implement an energy surplus/deficit feedback mechanism that disables cloning if there is an excess of energy and enables cloning if energy levels drop below the neuron’s requirements (see Fig. 4). The disable/enable signal applies to both mtDNA_*wild*_ and mtDNA_*del*_ even though mtDNA_*del*_ do not generate energy.

**Figure 4:**
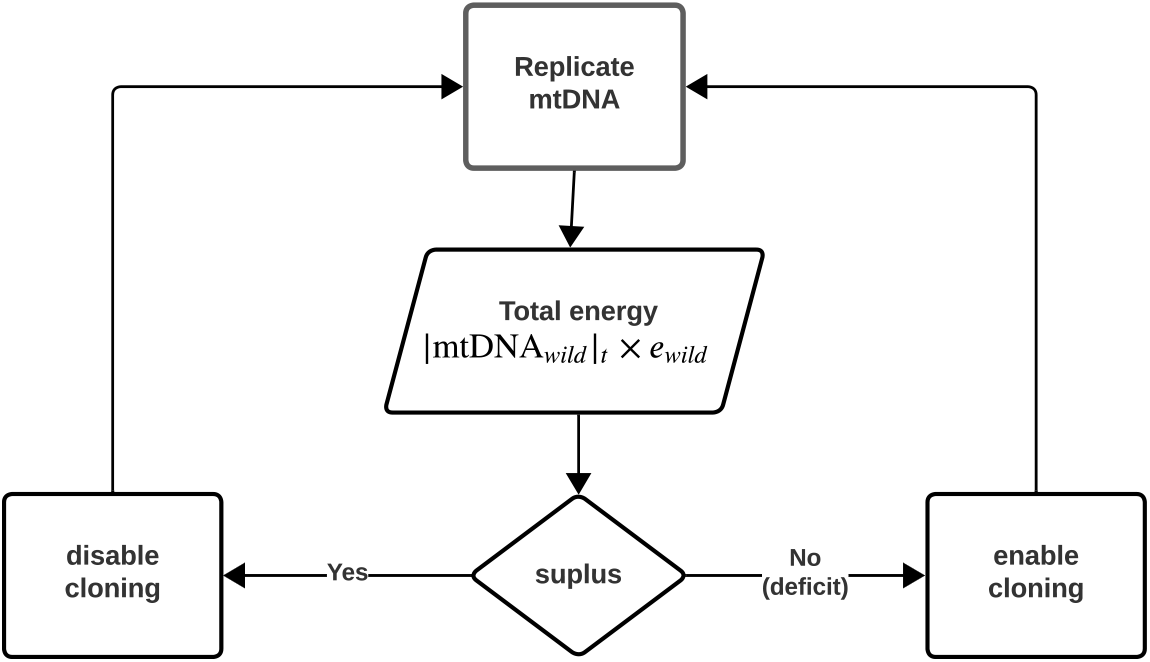
Energy surplus/deficit feedback mechanism. Given *e*_*wild*_ is the per-mtDNA_*wild*_ energy production and | mtDNA_*wild*_ |_*t*_ is the number of wild-type mtDNA at time *t*, if |mtDNA_*wild*_ |_*t*_ *× e*_*wild*_ is less than the cell’s energy demand (deficit) cloning is turned on (and off otherwise). Thus, the cell maintains a level of *CN*_*wild*_ (on average) for a given value of *e*_*wild*_, provided the organelle is not in competition. A change in *e*_*wild*_ invokes a change in *CN*_*wild*_ through the surplus/deficit feedback mechanism.

The neuron’s demand for energy is modelled as a consumer/producer mechanism. An mtDNA_*wild*_ produces *e*_*wild*_ units of energy while the neuron consumes (or at least attempts to) *E*_*cell*_ units every *t*. Thus, for each *t* there is an energy level *E*(*t*), such that:

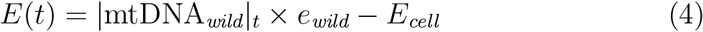

where |*X* |_*t*_ is the cardinality of *X* at time *t*. If *E*(*t*) ≥ *E_cell_*, then there is a surplus and a deficit otherwise.

We implement a capacity limit *C*_*max*_, whereby no mtDNA can replicate if:

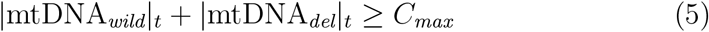

Initially (*t* → 0), there is an abundance of space within the organelle and there is spare capacity for all species of mtDNA. However, any clonal expansion may cause the aggregate population to reach *C*_*max*_ thereby disabling cloning. Replication is only (re)enabled after some attrition. The organelle oscillates between off/on periods of replication and mtDNA species enter a competition for space. We define the moment that the population reaches *C*_*max*_ for the first time as the *competition point*.

Cloning is a sequential process, that is, an mtDNA cannot simultaneously clone multiple times and must complete its current cloning process before entering the next. While undergoing cloning, mtDNA mutate according to the success of:

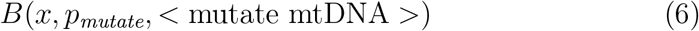

resulting in a new species of deletion mutant rather than a copy of its parent. In our simulation, a deletion mutant is half the size of its parent. Deletions may themselves mutate, leading to increasingly smaller molecules. The replication time is proportional to the mtDNA’s size, *S*_*m*_ (in nucleotides):

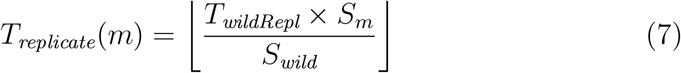

where *T*_*wildRepl*_ is the time for mtDNA_*wild*_ to replicate (2 hours) and *S*_*wild*_ is the size of a mtDNA_*wild*_ (16,569 nucleotides).

Table 1 shows the parameters of the model common to every simulation run.

**Table 1:** Parameters common to all runs of the simulation.

| Parameter | Value | Comment |
| --- | --- | --- |
| $t_{\text{max}}$ | 3,504,000 | Simulation run length (100 years) |
| $p_{\text{clone}}$ | $0.01 \text{ h}^{-1}$ | Clone probability |
| $T_{\text{wildRepl}}$ | 2 hours | Wild-type replication time |
| $TTL$ | 10 | Equiv. 30 day half-life |
| $p_{\text{decr}}$ | 0.0034 | |

## 4 Analysis

The primary function of mitochondria is oxidative phosphorylation, therefore, it is plausible that the decrease in the partial pressure of oxygen associated with an increase in altitude [27] would have an impact on mitochondrial function. A significant change (due to altitude) in the mtDNA copy number has been observed in several species, both mammalian and avian [21]. For all the species tested, the mtDNA copy number for animals at 3,000m was significantly reduced when compared to their counterparts below 500m. As this effect was observed across phyla, it suggests that it is a conserved adaptive response to a change in altitude. If the observed *in vitro* effects [18] translate to the *in vivo* situation then this adaptation could explain the reduced incidence of dementia associated with high altitude.

Any change to mtDNA_*wild*_ energy output efficiency causes a change in the level of *CN*_*wild*_ brought about by the energy feedback mechanism’s response to meet the cell’s energy demands. Thus *CN*_*wild*_ is a function of the cell’s energy demands *E*_*cell*_ and the per-mtDNA_*wild*_ energy production, *e*_*wild*_.

Mitochondrial oxidative phosphorylation is dependent on oxygen [34] and is sensitive to changes in pH and the partial pressure of oxygen. Lages *et al*. investigated how a change in the oxygen concentration altered mitochondrial function in human neural progenitor cells *in vitro*. Their results showed that the rates of oxidative phosphorylation of cells cultured for 18 days at an oxygen concentration of 0.21 and 0.03. At a low oxygen concentration the efficiency of oxidative phosphorylation increased, effectively increasing the oxygen flux and the P/O ratio.

We express changes in ATP production due to an increase in altitude as an *energy increase factor δ*_*x*_, where *x* is the oxygen concentration. That is, if 0.21 oxygen concentration is typical for sea level, this corresponds to a *baseline* ATP generation of 14.77 *pmol s*^−1^ *×* 10^−6^cells and, at this level, *δ*_0.21_ = *×* 1.0. At 0.03 oxygen concentration, ATP generation is equivalent to 26.41 *pmol s*^1^ 10^−6^cells and, thus, *δ*_0.03_ = 1.8. Assuming the energy increase factor *δ*_*x*_ changes linearly with oxygen concentration *x*, the straight line joining the two data points, namely (0.21, 1.01) and (0.03, 1.8), yields the expression:

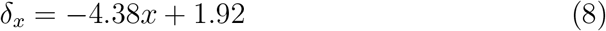

At 4,000m, a decrease of ≈ 20% has been observed in the mitochondrial volume in muscle cells compared to sea level [22, 15, 16]. We assume a similar effect in neurons. We further assume that any decrease in mitochondrial volume yields a proportional reduction in *C*_*max*_ (the mtDNA capacity of our simulated mitochondria). As we set *C*_*max*_ = 10, 000 at sea level then *C*_*max*_ = 8, 000 at an altitude of 4,000m. A straight line between these two point gives the relationship:

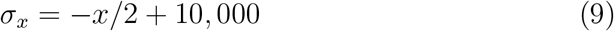

*C*_*max*_ is given by the function *σ*_*x*_ where *x* is the altitude (in meters). Table 2 shows the values of *CN*_*wild*_ and *C*_*max*_ for altitudes 1,600 and 3,000m, respectively.

**Table 2:** Parameter values for living at sea level and altitude with associated predisposition.

| Key | Altitude (m) | $CN_{wild}$ | $C_{max}$ | $p_m h^{-1}$ |
| --- | --- | --- | --- | --- |
| H/S | 0 | 1,000 | 10,000 | $2 \times 10^{-4}$ |
| H/M | 1,600 | 854 | 9,200 | $2 \times 10^{-4}$ |
| H/H | 3,000 | 764 | 8,500 | $2 \times 10^{-4}$ |
| L/S | 0 | 1,000 | 10,000 | $1 \times 10^{-4}$ |

We also consider high and low predisposition to dementia by modulating the mutation probability, where *p*_*m*_ = 2 *×* 10^−4^ *h*^−1^ and *p*_*m*_ = 1 *×* 10^−4^ *h*^−1^ are the respective mutation probabilities. We run 200 replications of four experiments:

- High predisposition at sea level (key: H/S),
- High predisposition at medium altitude (key: H/M),
- High predisposition at high altitude (key: H/H),
- Low predisposition at sea level (key: L/S).

Note that, *CN*_*wild*_ is a not actually a parameter that we set directly in the simulation, rather it is function (due to the energy feedback mechanism) of *e*_*wild*_, which we *do* set (adjusted according to *δ*_*x*_). However, for convenience, we refer to *CN*_*wild*_ as a parameter as it is, albeit indirectly, a factor of altitude and one that has been observed to change with altitude [21].

The graph in Fig. 5 shows the neuron loss rates for each experiment, H/S, H/M, H/H and L/S. It shows that for hosts with a high predisposition to dementia, living at altitude (H/M and H/H), the severity of dementia is reduced compared to living at sea level. Living at altitude pushes onset of dementia to age 65 and 67 for H/M and H/H, respectively; compared to H/S where onset is 59. Clearly living at altitude will not reduce severity and onset to low predisposition levels (onset of 87 years for L/S), however, there are some moderate benefits.

**Figure 5:**
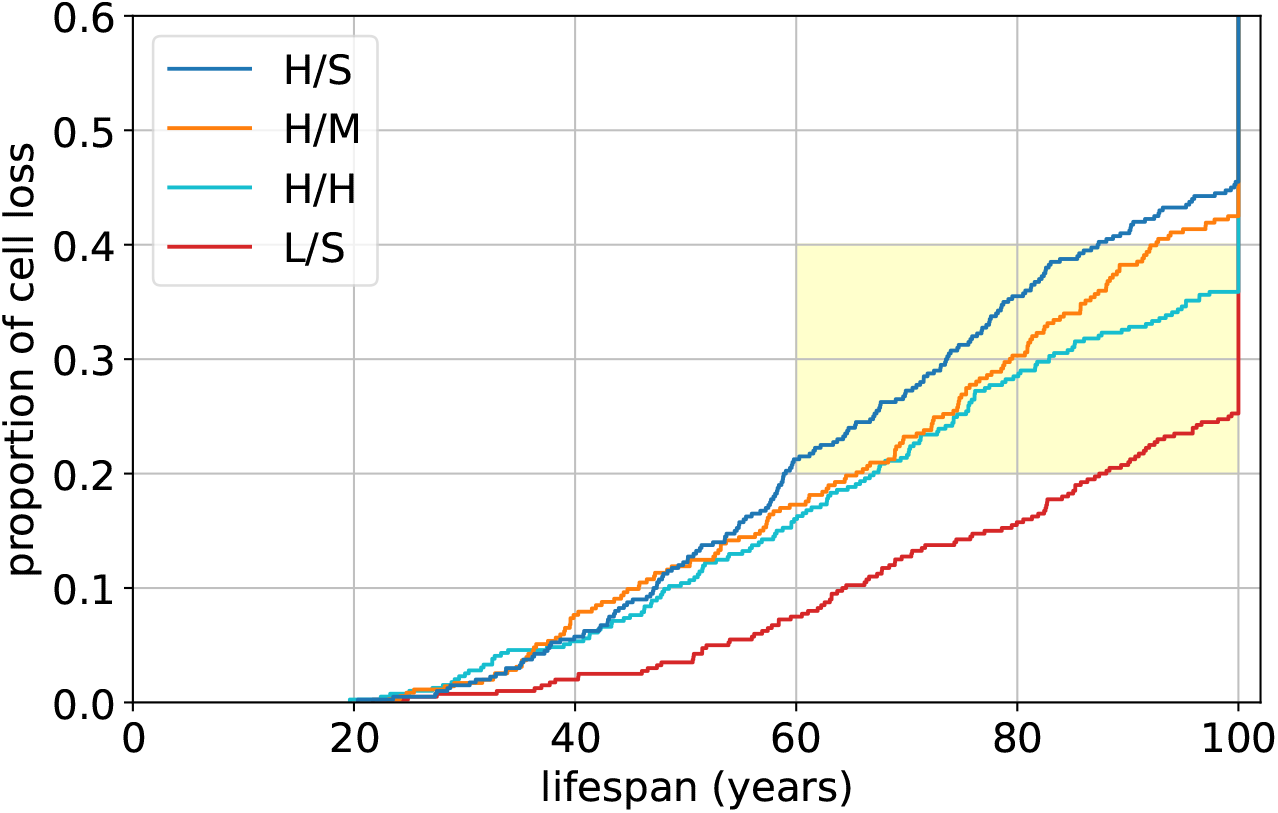
Neuron loss rates for a host at sea level versus altitude. The graph shows results for predisposition (high and low) at sea level in order to establish baselines. It also shows loss rates for hosts with high predisposition living at medium and high altitude. The yellow shaded area highlights the 20-40% neuron loss *dementia* region for the elderly.

From the data we can examine mtDNA dynamics within the organelle. The strip-plot in Fig. 6 (top) shows the heteroplasmy (number of mutant species) for both survived and deceased cells. It shows that heteroplasmy is greater for deceased cells than survived, which is to be expected as survived cells are not, typically, overrun by large mutant populations and rarely hit the competition point. A small proportion of survived cells do exhibit high heteroplasmy. Here, there would have been a build up of mutants and, while the neuron would have been in competition, it survived until the end of the simulation. However, the important result here is that there is a reduction, albeit moderate, in heteroplasmy with higher altitude.

**Figure 6:**
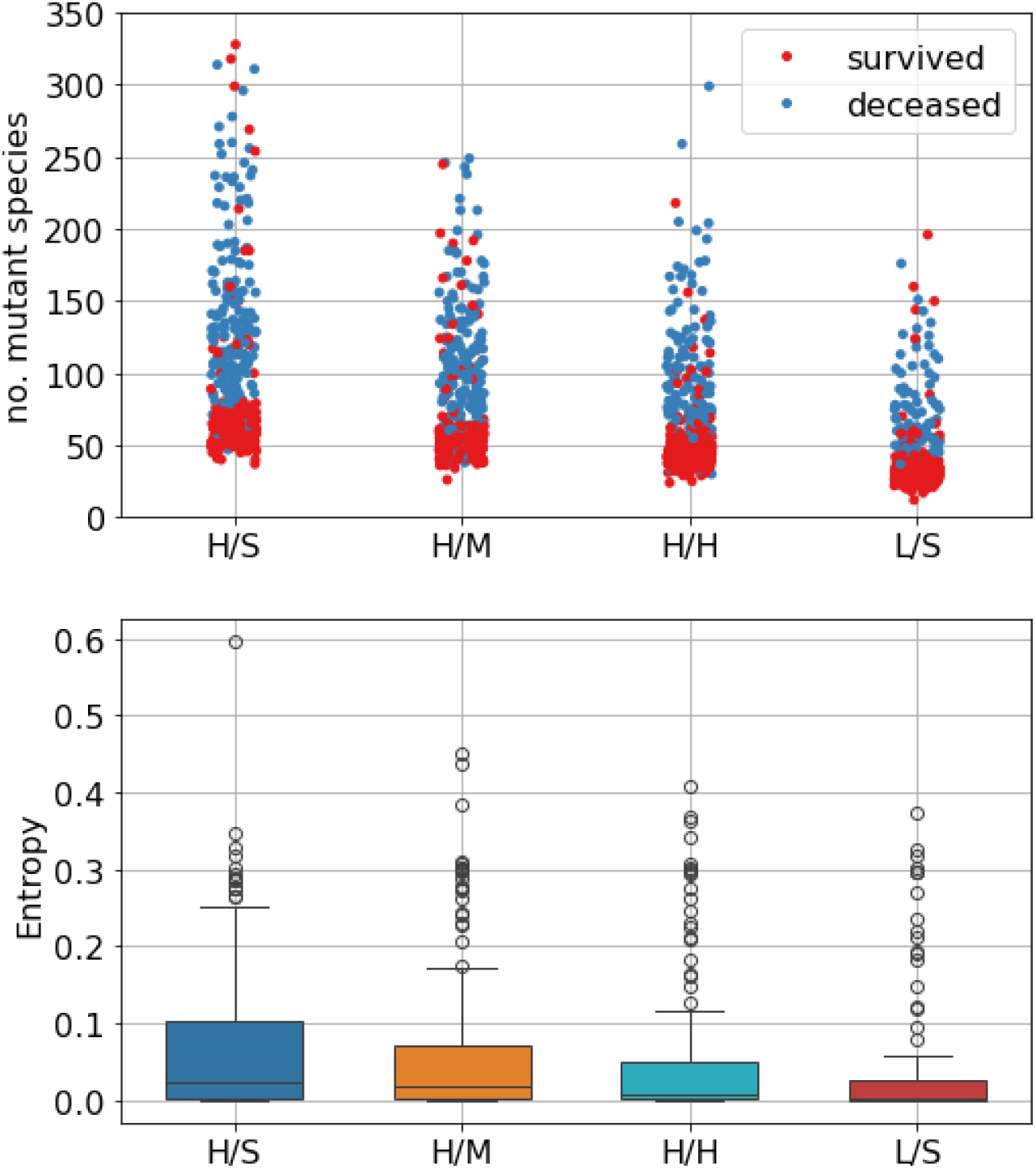
(Top) Heteroplasmy expressed as the number of mutant species that existed during a host’s lifetime. (Bottom) Entropy - a measure of the evenness of the distribution of mtDNA_*del*_ across species (deceased cells). The lower the number the greater the mutant population is dominated by clonal expansions.

In addition to heteroplasmy we are also interested in the nature of clonal expansion. In the case of deceased cells there are, typically, 1-4 clonal expansions and a small background of heteroplasmy [13], which is consistent with published results [5, 24]. We take a snapshot of the mutant population at the competition point and apply information entropy [29] to get a measure of evenness of the distribution of the number of mtDNA_*del*_ across the various species. The distribution of mutant species tends towards evenness at high entropy values an unevenness at low. The number of mutants in the *i*^*th*^ species is *n*_*i*_ and *p*_*i*_ is the normalised number of mutants. Entropy is, then, given by:

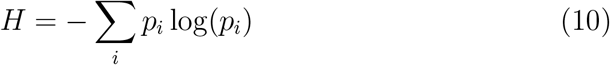

where:

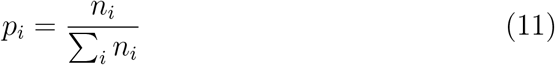

We exclude survived cells from the analysis as they tend to be devoid of large mutation populations, thus, we apply entropy to deceased cells only. From the graph in Fig. 6 (bottom) we can see that entropy decreases with altitude, thus, the population is dominated by fewer (but larger) clonal expansions. However, we should note, the effect is small.

Figure 7 shows the correlation between competition point and lifespan for deceased cells (survived cells rarely hit the competition point), in that, the earlier the competition point occurs, the shorter the neuron’s lifespan. However, neither predisposition or altitude have much effect on this correlation.

**Figure 7:**
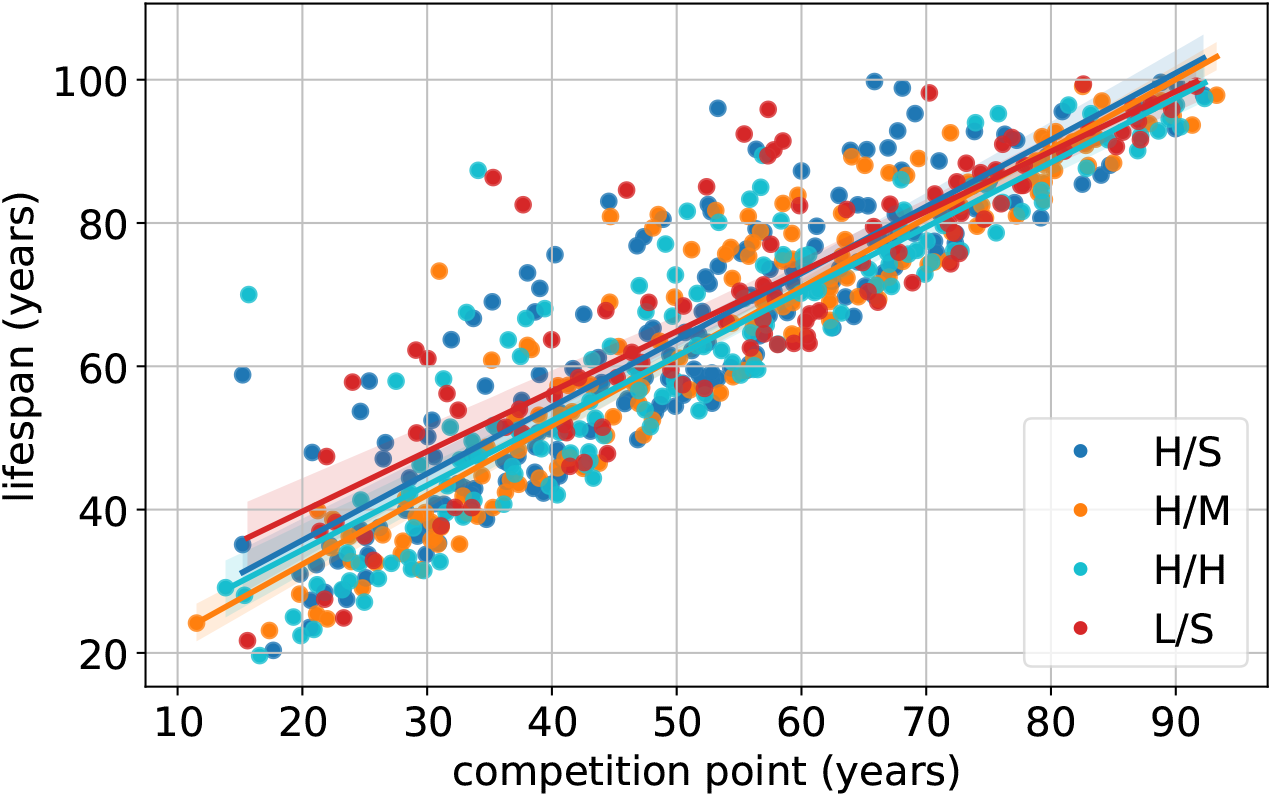
Lifespan versus competition point of deceased cells.

## 5 Discussion

In this paper we examined the effect of altitude on cognitive decline in humans with regard to mitochondrial dysfunction. Our aim was to provide an explanation as to why living at altitude affects the prevalence of dementia. We simulated proliferation of mtDNA_*del*_ in neurons where clonal expansion could interfere with the ability of mtDNA_*wild*_ to meet a neuron’s energy requirements, resulting in its premature demise. We recorded the neuron loss rates over a human lifespan of 100 years. We equate neuron loss to cognitive decline and we deem 20% loss to be the onset of dementia and that severity increases as losses exceed this threshold. While we ran each simulation for the full 100 years, it is unlikely that an actual host would survive beyond losses of 40%.

Using our simulator, we ran two experiments at sea level for high and low predisposition which we assign keys H/S and L/S, respectively. Two further experiments were performed for a host with high predisposition at medium and high altitude (keys: H/M and H/H, respectively). We can see from the results the most significant factor is predisposition. Predisposition to dementia is governed by mutation probability where onset of dementia was 59 and 87 years for high (H/S) and low predisposition (L/S), respectively. However, this result was not particularly surprising and is something we have demonstrated in previous publications [13]. These sea level results merely serve as baselines for neuron loss where H/S, in particular, is our control. For a host with high predisposition, living at altitude (H/M and H/H) reduced severity of dementia and delayed onset by several years compared to sea level (H/S). However, the benefits were moderate compared to low predisposition living at sea level (L/S).

The physiological changes due to altitude are predominantly due to the decrease in the partial pressure of oxygen and is associated with a decrease in copy number and mitochondrial volume. Thus, in our simulation of mitochondria, altitude affected *CN*_*wild*_ and *C*_*max*_. Increasing altitude reduces oxygen partial pressure which, in turn, increases per-mtDNA_*wild*_ energy production. Given we operate an energy surplus/deficit feedback mechanism to regulate mtDNA_*wild*_, an increase in per-mtDNA_*wild*_ energy production would down-regulate *CN*_*wild*_. We used *in vitro* results of ATP changes with oxygen concentration from Lages *et al*. in order establish the scale of per-mtDNA_*wild*_ energy production increase.

Changes in mitochondria volume are also associated with living at altitude. We equate changes in volume with the mtDNA capacity limit, specifically *C*_*max*_ diminshes (with volume) as altitude increases.

Lowering both these parameters, as altitude does, would have had opposing effects on neuron loss. We know from previous work that reducing *CN*_*wild*_ from low thousands to mid/high hundreds reduces incidence of dementia in the elderly [14]. Whereas, lowering *C*_*max*_ would have the effect of bringing the competition point forward because the mtDNA capacity is lower and any clonal expansion would reach a lower *C*_*max*_ earlier than a higher one (pursuant to living at sea level). It can be seen from Fig. 7 that competition point and lifespan are directly correlated, thus, the earlier a neuron reaches competition point the lower its life expectancy.

Overall, there is a moderate benefit to living at altitude versus sea level with regard to dementia in the elderly. It is interesting to note that the difference is, indeed, in the elderly and that there is little difference in cognitive decline in the young and middle aged. Furthermore, there is no noticeable difference between medium and high altitudes until a host reaches their 80s.

In our preliminary analysis in Section 2 we collected data sets from various regions, namely, European countries, Chinese provinces and US counties. These results indicate that living at altitude yields a moderate decrease in the incidence of dementia. This trend is most evident in the US and China data as there are large centers of population located over a wide range of altitudes. However, the trend is less apparent in Europe as it lacks the range of population centers of sufficient size at higher altitudes.

A confounding variable is the standardisation of the diagnostic criteria across countries as has been previously noted in Urrunaga-Pastor *et al*. [31]. We have no assurance that different European countries operate to the same diagnostic standard. However, the publications that accompany the US and Chinese data sets claim to adopt the DSM.

We need to stress we are not proposing *living at altitude* as a therapy for dementia. For one, it only benefits those with a high predisposition and it is not possible (at this time) to know a host’s predisposition ahead of onset. Furthermore, it is not sustainable for a large proportion of the world’s population to relocate to high altitudes if they were to second guess their predisposition. An examination of a number data sets on regional dementia leads us to expect a lower, albeit moderate, prevalence of dementia with habitation at higher altitudes.

We used simulation techniques in order to provide a possible explanation for this phenomenon. We acknowledge that dementia is a complex condition and likely influenced by multiple factors. However, the focus of this paper was the effects of living at altitude on mitochondrial dysfunction brought about by clonal expansion of deletion mutants. The results of our simulation show that the effects of living at altitude lead to lower levels of neuron loss and are consistent with dementia prevalence data sets when plotted against altitude.

## 6 Conclusions

In this paper we have studied the effects on mitochondrial dysfunction that living at altitude has on the prevalence of dementia in the elderly. We used simulation methods to model proliferation of mtDNA deletion mutations and the resulting neuron loss.

It has been reported that living at altitude reduces copy number and mitochondrial volume, thus, we set these parameters in our pseudo mitochondrial organelle in order to simulate a host with high predisposition to dementia living at medium and high altitudes where our control was a host living at sea level.

Regional data sets show that there is some evidence of a link between living at altitude and prevalence of dementia. This link aligns with our simulation results which show that living at altitude pushes back onset of dementia by a few years and yields a modest reduction in severity. Furthermore, the benefits to living at altitude are in the elderly; there does not appear to be any effect on cognitive decline in the young and middle aged.

A mechanism or therapeutic intervention that mimics the effects of living at altitude, that is, reduces the P/O ratio similar to the effects of low oxygen tension, could be an effective treatment for dementia.

Aging is the main risk factor for various pathologies and neurodegenerative conditions. The world has an aging population where the percentage of over 65s is already more than 20% in some countries. Even a modest delay in the onset of dementia or a reduction in the incidence rate would have a profound social and financial benefit.

## Appendix

### Algorithm 1

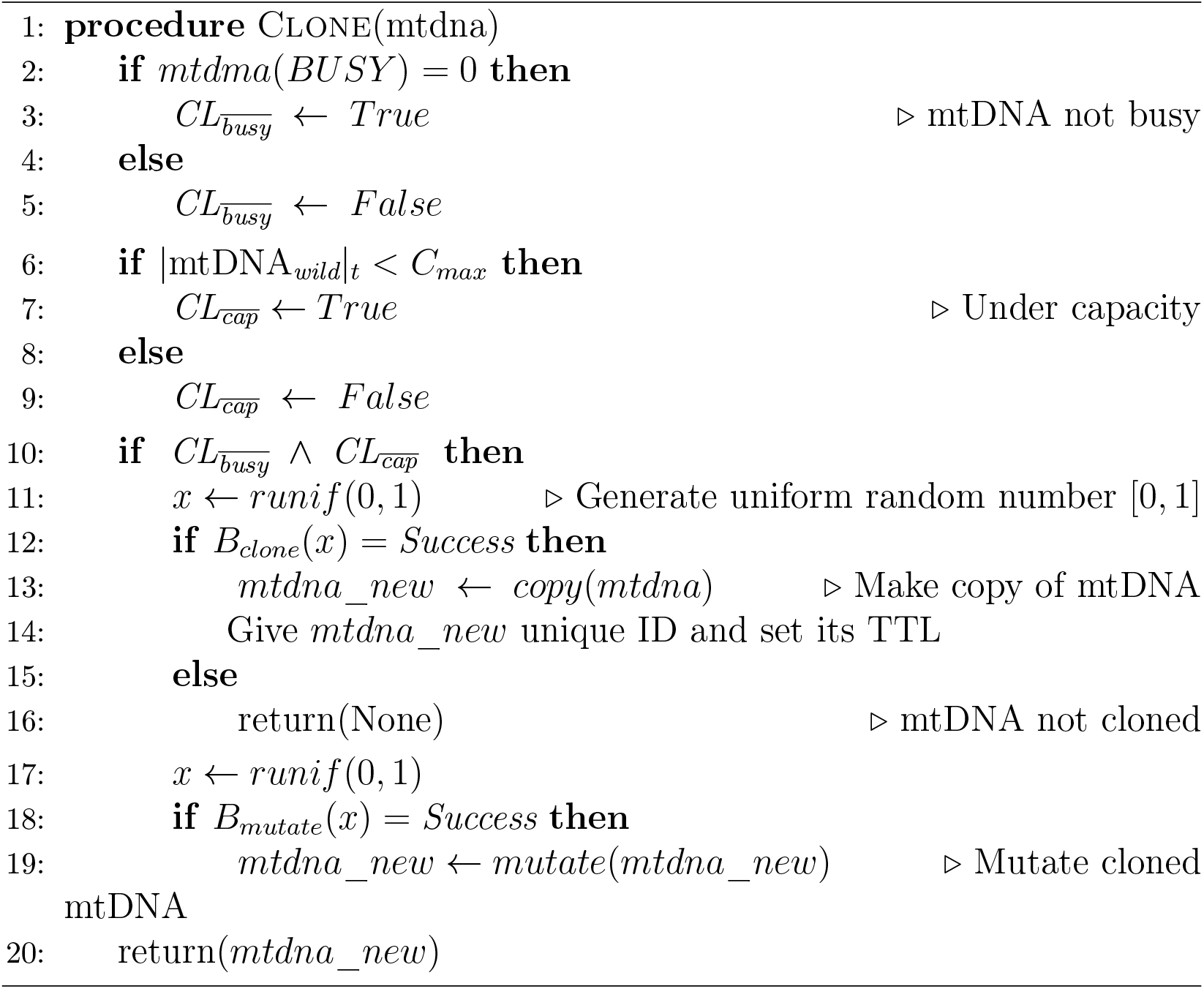

Simulator

### Algorithm 2

Simulator

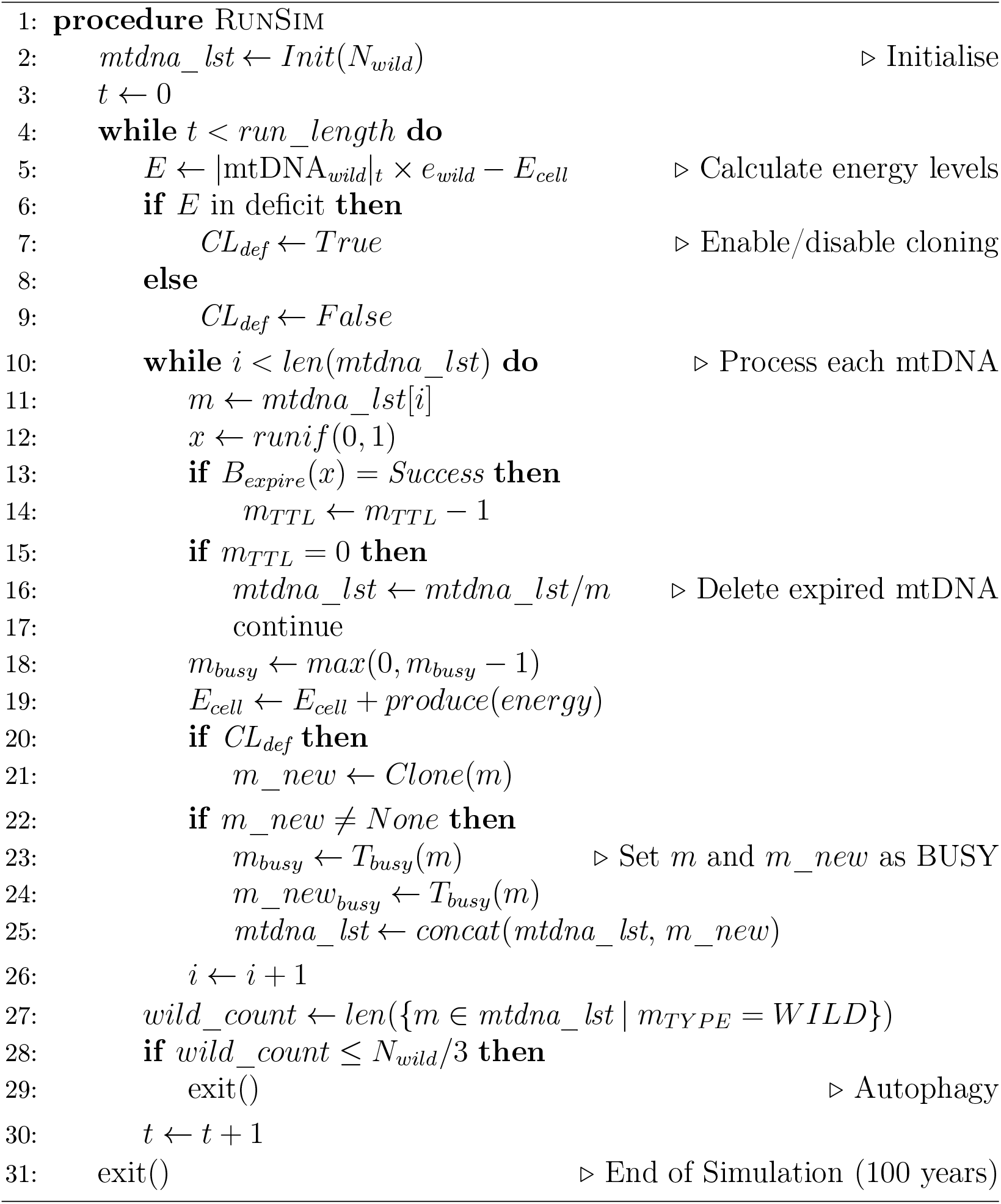

## References

[1] Alzheimer Europe Foundation. Prevalence of dementia in europe. https://www.alzheimer-europe.org/{\protect\penalty\z@}dementia/{\protect\penalty\z@}prevalence-dementia-{\protect\penalty\z@}europe, 2023.

[2] C H Andrade-Moraes, A V Oliveira-Pinto, E Castro-Fonseca, C G da Silva, D M Guimaraes, D Szczupak, D R Parente-Bruno, L R B Carvalho, L Polichiso, B V Gomes, L M Oliveira, R D Rodriguez, R E P Leite, R E L Ferretti-Rebustini, W Jacob-Filho, C A Pasqualucci, T T Grinberg, and R Lent. Cell number changes in Alzheimer’s disease relate to dementia, not to plaques and tangles. Brain, 136(12):3738–3752, 2013.

[3] Thomas Arendt, Martina K Bruckner, Markus Morawski, Carsten Jager, and Hermann-Josef Gertz. Early neurone loss in alzheimer’s disease: cortical or subcortical? Acta neuropathologica communications, 3(1):1– 11, 2015.

[4] Ben Sundra Ashok, Thekkuttuparambil Ananthanarayanan Ajith, and Senthilkumar Sivanesan. Hypoxia-inducible factors as neuroprotective agent in alzheimer’s disease. Clinical and Experimental Pharmacology and Physiology, 44(3):327–334, 2017.

[5] Tiziana Casoli, Liana Spazzafumo, Giuseppina Di Stefano, and Fiorenzo Conti. Role of diffuse low-level heteroplasmy of mitochondrial DNA in Alzheimer’s disease neurodegeneration. Frontiers in Aging Neuroscience, 7:4106, 2015.

[6] Klodian Dhana, Todd Beck, Pankaja Desai, Robert S Wilson, Denis A Evans, and Kumar B Rajan. Prevalence of alzheimer’s disease dementia in the 50 us states and 3142 counties: A population estimate using the 2020 bridged-race postcensal from the national center for health statistics. Alzheimer’s & Dementia, 2023.

[7] DL Dickstein, D Kabaso, AB Rocher, JI Luebke, SL Wearne, and PR Hof. Changes in the structural complexity of the aged brain. Aging Cell, 6(3):275–284, 2007.

[8] Ligia J Dominguez, Nicola Veronese, Laura Vernuccio, Giuseppina Catanese, Flora Inzerillo, Giuseppe Salemi, and Mario Barbagallo. Nutrition, physical activity, and other lifestyle factors in the prevention of cognitive decline and dementia. Nutrients, 13(11):4080, 2021.

[9] Lei Dong, Rui Du, and Yu Liu. Mapping evolving population geography in china: Spatial redistribution, regional disparity, and urban sprawl. Regional Disparity, and Urban Sprawl (March 4, 2022), 2022.

[10] Josef Finsterer. Cognitive decline as a manifestation of mitochondrial disorders (mitochondrial dementia). Journal of the neurological sciences, 272(1-2):20–33, 2008.

[11] AF Fotenos, AZ Snyder, L. Girton, JC Morris, and RL Buckner. Normative estimates of cross-sectional and longitudinal brain volume decline in aging and AD. Neurology, 64(6):1032–9, 2005.

[12] Catherine M Hill, Dagmara Dimitriou, Ana Baya, Rebecca Webster, Johanna Gavlak-Dingle, Veline Lesperance, Kate Heathcote, and Romola S Bucks. Cognitive performance in high-altitude andean residents compared with low-altitude populations: from childhood to older age. Neuropsychology, 28(5):752, 2014.

[13] Alan G Holt and Adrian M Davies. A comparison of mtdna deletion mutant proliferation mechanisms. Journal of Theoretical Biology, 551:111244, 2022.

[14] Alan G Holt and Adrian M Davies. The prevalence of dementia in humans could be the result of a functional adaptation. Computational Biology and Chemistry, 106:107939, 2023.

[15] Hans Hoppeler and Michael Vogt. Muscle tissue adaptations to hypoxia. Journal of experimental biology, 204(18):3133–3139, 2001.

[16] James A Horscroft, Aleksandra O Kotwica, Verena Laner, James A West, Philip J Hennis, Denny ZH Levett, David J Howard, Bernadette O Fernandez, Sarah L Burgess, Zsuzsanna Ament, et al. Metabolic basis to sherpa altitude adaptation. Proceedings of the National Academy of Sciences, 114(24):6382–6387, 2017.

[17] Yume Imahori, Davide L Vetrano, Xin Xia, Giulia Grande, Petter Ljungman, Laura Fratiglioni, and Chengxuan Qiu. Association of resting heart rate with cognitive decline and dementia in older adults: A population-based cohort study. Alzheimer’s & Dementia, 18(10):1779–1787, 2022.

[18] Yury M Lages, Juliana M Nascimento, Gabriela A Lemos, Antonio Galina, Leda R Castilho, and Stevens K Rehen. Low oxygen alters mitochondrial function and response to oxidative stress in human neural progenitor cells. PeerJ, 3:e1486, 2015.

[19] Jerzy Leszek, Elizaveta V Mikhaylenko, Dmitrii M Belousov, Efrosini Koutsouraki, Katarzyna Szczechowiak, Małgorzata Kobusiak-Prokopowicz, Andrzej Mysiak, Breno S Diniz, Siva G Somasundaram, Cecil E Kirkland, et al. The links between cardiovascular diseases and alzheimer’s disease. Current Neuropharmacology, 19(2):152–169, 2021.

[20] Yuan Li and Yan Wang. Effects of long-term exposure to high altitude hypoxia on cognitive function and its mechanism: A narrative review. Brain Sciences, 12(6):808, 2022.

[21] Rui Liu, Long Jin, Keren Long, Qianzi Tang, Jideng Ma, Xun Wang, Li Zhu, An’an Jiang, Guoqing Tang, Yanzhi Jiang, et al. Analysis of mitochondrial dna sequence and copy number variation across five high-altitude species and their low-altitude relatives. Mitochondrial DNA Part B, 3(2):847–851, 2018.

[22] Andrew J Murray and James A Horscroft. Mitochondrial function at extreme high altitude. The Journal of physiology, 594(5):1137–1149, 2016.

[23] Li nan Cheng, Li Zhao, Xiao feng Xie, Liang Wang, Xiu ying Hu, Xiao yang Dong, and Feng ying Zhang. Care willingness and demand of residents under 60 years of age in western china: a cross-sectional study. BMJ open, 11(8), 2021.

[24] Gonzalo S Nido, Christian Dölle, Irene Flønes, Helen A Tuppen, Guido Alves, Ole-Bjørn Tysnes, Kristoffer Haugarvoll, and Charalampos Tzoulis. Ultradeep mapping of neuronal mitochondrial deletions in Parkinson’s disease. Neurobiology of aging, 63:120–127, 2018.

[25] Katie A O’Brien, Tatum S Simonson, and Andrew J Murray. Metabolic adaptation to high altitude. Current Opinion in Endocrine and Metabolic Research, 11:33–41, 2020.

[26] Jiwon Park, Sunhee Jung, Sang-Min Kim, In young Park, Ngan An Bui, Geum-Sook Hwang, and Inn-Oc Han. Repeated hypoxia exposure induces cognitive dysfunction, brain inflammation, and amyloidβ/p-tau accumulation through reduced brain o-glcnacylation in zebrafish. Journal of Cerebral Blood Flow & Metabolism, 41(11):3111–3126, 2021.

[27] Andrew J Peacock. Oxygen at high altitude. Bmj, 317(7165):1063–1066, 1998.

[28] Chengxuan Qiu, Bengt Winblad, and Laura Fratiglioni. The age-dependent relation of blood pressure to cognitive function and dementia. The Lancet Neurology, 4(8):487–499, 2005.

[29] CE Shannon and W Weaver. The Mathematical Theory of Communication. University of Illinois Press, Urbana IL, 1949.

[30] Stephen Thielke, Christopher G Slatore, and William A Banks. Association between alzheimer dementia mortality rate and altitude in california counties. JAMA psychiatry, 72(12):1253–1254, 2015.

[31] Diego Urrunaga-Pastor, Diego Chambergo-Michilot, Fernando M Runzer-Colmenares, Josmel Pacheco-Mendoza, and Vicente A Benites-Zapata. Prevalence of cognitive impairment and dementia in older adults living at high altitude: A systematic review and meta-analysis. Dementia and Geriatric Cognitive Disorders, 50(2):124–134, 2021.

[32] Vittore Verratti, Claudio Ferrante, Davide Soranna, Antonella Zambon, Suwas Bhandari, Giustino Orlando, Luigi Brunetti, and Gianfranco Parati. Effect of high-altitude trekking on blood pressure and on asymmetric dimethylarginine and isoprostane production: Results from a mount ararat expedition. The Journal of Clinical Hypertension, 22(8):1494–1503, 2020.

[33] Christopher S von Bartheld. Myths and truths about the cellular composition of the human brain: A review of influential concepts. Journal of Chemical Neuroanatomy, 93:2–15, 2018.

[34] David F Wilson, David K Harrison, and Sergei A Vinogradov. Oxygen, ph, and mitochondrial oxidative phosphorylation. Journal of Applied Physiology, 113(12):1838–1845, 2012.

[35] World topographic map. Topographic maps. https://en-us.topographic-map.com/, 2024.

[36] Yu-Tzu Wu, Hsin-yi Lee, Samuel Norton, Chuanfeng Chen, Hongxia Chen, Chenglin He, Jane Fleming, Fiona E Matthews, and Carol Brayne. Prevalence studies of dementia in mainland china, hong kong and taiwan: a systematic review and meta-analysis. PloS one, 8(6):e66252, 2013.

[37] Xinjuan Zhang and Jiaxing Zhang. The human brain in a high altitude natural environment: A review. Frontiers in Human Neuroscience, 16:915995, 2022.

